# SNP-sites: rapid efficient extraction of SNPs from multi-FASTA alignments

**DOI:** 10.1101/038190

**Authors:** Andrew J. Page, Ben Taylor, Aidan J. Delaney, Jorge Soares, Torsten Seemann, Jacqueline A. Keane, Simon R. Harris

## Abstract

Rapidly decreasing genome sequencing costs have led to a proportionate increase in the number of samples used in prokaryotic population studies. Extracting single nucleotide polymorphisms (SNPs) from a large whole genome alignment is now a routine task, but existing tools have failed to scale efficiently with the increased size of studies. These tools are slow, memory inefficient and are installed through non-standard procedures. We present *SNP-sites* which can rapidly extract SNPs from a multi-FASTA alignment using modest resources and can output results in multiple formats for downstream analysis. SNPs can be extracted from a 8.3 GB alignment file (1,842 taxa, 22,618 sites) in 267 seconds using 59 MB of RAM and 1 CPU core, making it feasible to run on modest computers. It is easy to install through the Debian and Homebrew package managers, and has been successfully tested on more than 20 operating systems. *SNP-sites* is implemented in C and is available under the open source license GNU GPL version 3.

## DATA SUMMARY

1. The source code for *SNP-sites* is available from GitHub under GNU GPL v3; (URL — https://github.com/sanger-pathogens/snp_sites)

2. *S. Typhi* multi FASTA alignment data has been deposited in Figshare: https://dx.doi.org/10.6084/m9.figshare.2067249.v1

We confirm all supporting data, code and protocols have been provided within the article or through supplementary data files.

## IMPACT STATEMENT

Rapidly extracting SNPs from increasingly large alignments, both in number of sites and number of taxa, is a problem that current tools struggle to deal with efficiently. *SNP-sites* was created with these challenges in mind and this paper demonstrates that it scales well, using modest desktop computers, to sample sizes far in excess of what is currently analysed in single population studies. The software has also been packaged to allow it to be easily installed on a wide variety of operating systems and hardware, something often neglected in bioinformatics.

## INTRODUCTION

As the cost of sequencing has rapidly decreased, the number of samples sequenced within a study has proportionately increased and now stands in the thousands (Chewapreecha *et al.*, 2014; Nasser *et al.*, 2014; Wong *et al.*, 2015). A common task in prokaryotic bioinformatics analysis is the extraction of all single nucleotide polymorphisms (SNPs) from a multiple FASTA alignment. Whilst it is a simple problem to describe, current tools cannot rapidly or efficiently extract SNPs in the increasingly large data sets found in prokaryotic population studies. These inefficiencies, such as loading all the data into memory (Lindenbaum, 2015), or slow speed due to algorithm design (Capella-Gutiérrez *et al.*, 2009), make it infeasible to analyse these sample sets on modest computers. Furthermore, existing tools employ challenging, non-standard installation procedures.

A number of applications exist which can extract SNPs from a multi FASTA alignment, such as JVarKit (Lindenbaum, 2015), TrimAl (Capella-Gutiérrez *et al.*, 2009), PGDSpider (Lischer and Excoffier, 2012) and PAUP⋆ (Swofford, 2002).

JVarKit is a Java toolkit which can output SNP positions in VCF format (Danecek *et al.*, 2011). The standardised VCF format allows for post-processing with BCFtools (Danecek *et al.*, 2011), which is used to analyse variation in very large datasets such as the Human 1000 Genomes project (Sudmant *et al.*, 2015). It is reasonably fast, however it uses nearly 8 bytes of RAM per base of sequencing, which results in substantial memory usage for even small data sets. For example a 1 GB alignment (200 taxa, 50,000 sites, 5 MBp genomes) required 7.2 GB of RAM. TrimAl (version 1.4) is a C++ tool which outputs variation, given a multiple FASTA alignment, however it does not support VCF format, only outputting the positions of SNPs in a bespoke format. It is very slow for small sample sets, however it uses less memory than JVarKit. PGDSpider is a Java based application which can output a VCF file, however the authors warn it is not suitable for large files, so it has been excluded from this analysis. PAUP⋆ is a popular commercial application but as it is no longer distributed it was not available for comparison. None of these applications are easily installable on a wide variety of operating systems and environments. TrimAl is the only application available in Flomebrew and none are available through the Debían package management system.

Fiere we present *SNP-sites* which overcomes these limitations by managing disk I/O and memory carefully, and optimizing the implementation using C (ISO C99 compliant). Standard installation methods are used, with the software prepackaged and available through the Debían and Flomebrew package managers. The software has been successfully run on more than 20 architectures using Debían Linux, Redhat Enterprise Linux and on multiple versions of OS X. A Cython version of the *SNP-sites* algorithm called PySnpSites (https://github.com/bewt85/PySnpSites) is also presented for comparison purposes.

## THEORY AND IMPLEMENTATION

The input to the software is a single multiple FASTA alignment of nucleotides, where all sequences are the same length and have already been aligned. The file can optionally be gzipped. This alignment may have been generated by overlaying SNPs on a consensus reference genome, or using a multiple alignment tool, such as MUSCLE (Edgar, 2004), PRANK (Löytynoja, 2014), MAFFT (Katoh and Standley, 2013), or ClustalW (Thompson *et al.*, 2002).

By default the output format is a multiple FASTA alignment. The output format can optionally be changed to PHYLIP format (Felsenstein, 1989) or VCF format (version 4.1) (Danecek *et al.*, 2011). When used as a preprocessing step for FastTree (Price *et al.*, 2010), this substantially decreases the memory usage of FastTree during phylogenetic tree construction. The PHYLIP format can be used as input to RAxML (Stamatakis, 2014) for creating phylogenetic trees. For phylogenetic reconstructions removing monomorphic sites from an alignment may require a different model to avoid parameters being incorrectly estimated. The VCF output retains the position of the SNPs in each sample and can be parsed using standard tools such as BCFtools (Danecek *et al.*, 2011) or for GWAS analysis using PUNK (Chang *et al.,* 2015).

Each sequence is read in sequentially. A consensus sequence is generated in the first pass and is iteratively compared to each sequence. The position of any difference is noted. A second pass of the input file extracts the bases at each SNP site and outputs them in the chosen format. Where a base is unknown or is a gap (n/N/?/-), the base is regarded as a non-variant.

For example, given the input alignment:

~~~
>sample1
AGACACAGTCAC
>sample2
AGACAC----AC
>sample3
AAACGCATTCAN
~~~

the output is:

~~~
>sample1
GAG
>sample2
GA-
>sample3
AGT
~~~

The maximum resource requirements of the algorithm are known. Given the number of SNP sites is p, the number of samples is s and the number of bases in a single alignment is g the maximum memory usage can be defined as:

max(p^×^s, g^×^2).

Given that f is the size of the input file and o is the size of the output file, the file I/O is defined as:

2^×^f< = I/O <= 2^×^f+o.

The computational complexity is O(n). These properties make the algorithm theoretically scalable and feasible on large datasets far beyond what is currently analysed within a single study.

All changes to *SNP-sites* are validated automatically against a hand generated set of example cases incorporated into unit tests. A continuous integration system (https://travis-ci.org/sanger-pathogens/snp_sites) ensures that modifications which change the output erroneously are publically flagged.

To test the performance of *SNP-sites,* we have compared it with JVarKit, TrimAl and PySnpSites (https://github.com/bewt85/PySnpSites). PySnpSites is a Cython based partial reimplementation of the *SNP-sites* algorithm. A number of simulated datasets were generated to exercise the different parameters and to see their effect on memory usage and running time. All of the software to generate these datasets is contained within the *SNP-sites* source code repository.

All experiments were performed using a single processor (2.1 Ghz AMD Opteron 6272) with a maximum of 16 GB of RAM available. The maximum run time of an application was set as 12 hours, after which time the experiment was halted.

Alignments were generated with varying numbers of SNPs to show the effect of SNP density on the performance of each application. Each alignment had 1,000 samples and a genome alignment length of 5 Mbp, with a total file size of 4.8 GB. This is a scale encountered in recent studies (Wong *et al.*, 2015). As the SNP density increases, so does the running time and memory usage as seen in Fig. 1(a) and Fig. 1(b). The running time of both *SNP-sites* and PySnpSites is reasonable however the memory usage of PySnpSites rapidly exceeds the maximum allowed memory (16 GB). Where 20% of bases in the input alignment are SNPs, *SNP-sites* uses only uses 1 GB of RAM, or approximately 20% of the file input size, scaling with the volume of variation rather than the size of the input file. In all experiments JVarKit exceeded the maximum allowed memory and was halted. All experiments using TrimAl exceeded the maximum running time of 12 hours. As both of these applications did not successfully complete they are not present in the results.

**Figure 1.**
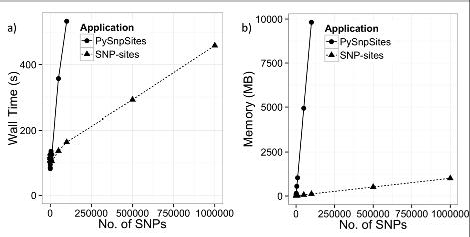
a) Wall time in seconds when the number of SNPs is varied, b) Memory in MB when the number of SNPs is varied. All JVarKit experiments exceeded the maximum memory and all TrimAl experiments exceeded the maximum run time, so are not shown

The number of samples analysed within a single study now stands in the thousands (Chewapreecha *et al.*, 2014). To cope with this scale and to demonstrate how applications will perform in the future, we generated alignments with 100 to 100,000 samples. Each genome contained 1 Mbp, and 1,000 SNP sites. The total file sizes ranged from 0.1 GB to 86 GB. As the number of taxa increase, the running time of PySnpSites and *SNP-sites* increases linearly, with *SNP-sites* taking 32 minutes to analyse an 86 GB alignment with 100,000 taxa as can be seen in Fig. 2(a). The running time of JVarKit is ten times greater than that of PySnpSites and *SNP-sites* as shown in Fig. 2(b), however it exceeds the 16 GB maximum memory limit beyond 1,000 taxa. The running time of TrimAl is another order of magnitude greater, making it rapidly infeasible to run. The memory usage of *SNP-sites* is substantially less than all other applications, with the closest, PySnpSites using 9.2 GB of RAM compared to 0.274 GB of RAM for *SNP-sites.*

**Figure 2.**
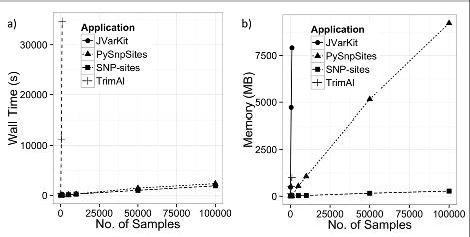
a) Wall time in seconds when the number of samples is varied, b) Memory in MB when the number of samples is varied.

Finally the length of each genome in the alignment is varied from 100,000 to 100 Mbp with 1,000 taxa and 1,000 SNP sites in each alignment. PySnpSites and *SNP-sites* performed consistently well as shown in Fig. 3(a), with both taking ≈40 minutes to process the largest 95 GB alignment file. *SNP-sites* uses just 203 MB of RAM compared to 691 MB by PySnpSites as shown in Fig. 3(b). The other two applications exceed the maximum running times and/or the maximum memory whilst trying to analyse 5 Mbp genomes, which is the size of a typical Gram negative bacterial genome.

**Figure 3.**
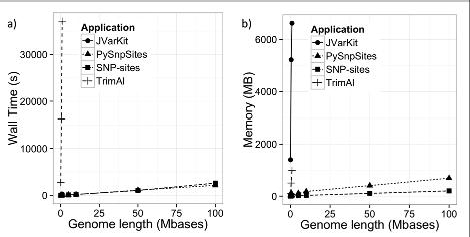
a) Wall time in seconds when the genome length is varied, b) Memory in MB when the genome length is varied.

The performance of *SNP-sites* was evaluated on a real data set of *Salmonella* Typhi from (Wong *et al.*, 2015). A total of 1,842 taxa were aligned to the 4.8 Mbp chromosome (accession number AL513382) of S. Typhi CT18. This gave a total alignment file size of 8.3 GB and incorporated SNPs at 22,618 sites. *SNP-sites* used 59 MB of RAM and took 267 seconds.

## CONCLUSION

Extracting variation from a multiple FASTA alignment is a common task, and whilst it is simple to define, existing tools fail to perform well. We showed that *SNP-sites* performed consistently under a variety of conditions, using low amounts of RAM and had a low running time for even for the largest datasets we simulated to represent the scale of studies expected in the near future. This makes it feasible to run on standard desktop machines. *SNP-sites* uses standard installation methods with the software prepackaged and available through the Debían and Homebrew package managers. The software has been successfully tested and run on more than 20 architectures using Debían Linux and on multiple versions of OS X.

## ACKNOWLEDGEMENTS

Thanks to Sascha Steinbiss, Andreas Tille and Nicholas J. Croucher for their assistance and advice. Thanks to Vanessa Wong and Kathryn Holt for providing the S. Typhi data set. This work was supported by the Wellcome Trust (grant WT 098051).

## ABBREVIATIONS

SNP: single nucleotide polymorphism
VCF: Variant Call Format

## DATA BIBLIOGRAPHY

1. Page, A.J., Github https://github.com/sanger-pathogens/snp_sites (2016).

2. Wong,V & Holt K., Figshare: https://dx.doi.org/10.6084/m9.figshare.2067249.vl (2016).

3. Parkhill, J., Genbank accession number AL513382 (2001).

